# Stable transfection in the protist *Corallochytrium limacisporum* allows identification of novel cellular features among unicellular relatives of animals

**DOI:** 10.1101/2020.11.12.379420

**Authors:** Aleksandra Kożyczkowska, Sebastián R. Najle, Eduard Ocaña-Pallarès, Cristina Aresté, Iñaki Ruiz-Trillo, Elena Casacuberta

## Abstract

The evolutionary path from protists to multicellular animals remains a mystery. Recent work on the genomes of several unicellular relatives of animals has shaped our understanding of the genetic changes that may have occurred in this transition. However, the specific cellular modifications that took place to accommodate these changes remain unclear. Functional approaches are now needed to unravel how different cell biological features evolved. Recent work has already established genetic tools in three of the four unicellular lineages closely related to animals (choanoflagellates, filastereans, and ichthyosporeans). However, there are no genetic tools available for Corallochytrea, the lineage that seems to have the widest mix of fungal and metazoan features, as well as a complex life cycle. Here, we describe the development of stable transfection in the corallochytrean *Corallochytrium limacisporum*. Using a battery of cassettes to tag key cellular components, such as nucleus, plasma membrane, cytoplasm and actin filaments, we employ live imaging to discern previously unknown biological features of *C. limacisporum*. In particular, we identify two different paths for cell division—binary fission and coenocytic growth—that reveal a non-linear life cycle in *C. limacisporum*. Additionally, we found that *C. limacisporum* is binucleate for most of its life cycle, and that, contrary to what happens in most eukaryotes, nuclear division is decoupled from cell division. The establishment of these tools in *C. limacisporum* fills an important gap in the unicellular relatives of animals, opening up new avenues of research with broad taxon sampling to elucidate the specific cellular changes that occurred in the evolution of animals.

## INTRODUCTION

Multicellular animals, or metazoans, display many characteristic cell biological features, such as an embryonic development and cell differentiation among others (Martin et al., 2008; Alberts et al., 2002). To understand how such features evolved, however, we need to compare metazoan cells with those of their extant relatives. Metazoa belong to the Holozoa clade, which also includes four different unicellular lineages: choanoflagellates, filastereans, ichthyosporeans, and corallochytreans/pluriformeans (Fig. 1A) (King, 2005; Shalchian-Tabrizi et al., 2008; Mendoza et al., 2002; Torruella et al., 2015; Hehenberger et al., 2017). These lineages display a wide range of morphologies, life stages and developmental patterns, some of them potentially homologous to animal ones. Interestingly, each of the unicellular holozoan clades contain species that can transiently form multicellular structures. For example, colony formation by clonal division is observed in some choanoflagellates (Fairclough et al., 2010; Dayel et al., 2011) an aggregative stage is present in some filastereans (Sebé-Pedrós et al., 2013) and coenocytic growth is found in some of the ichthyosporeans (Suga and Ruiz-Trillo, 2013) and corallochytreans (Tikhonenkov et al., 2020^a^) that have been described to date (Fig. 1B). Thus, comparative studies in Holozoa provide a unique opportunity to understand the evolution of different eukaryotic features, including the evolution of animal multicellularity.

**Figure 1.**
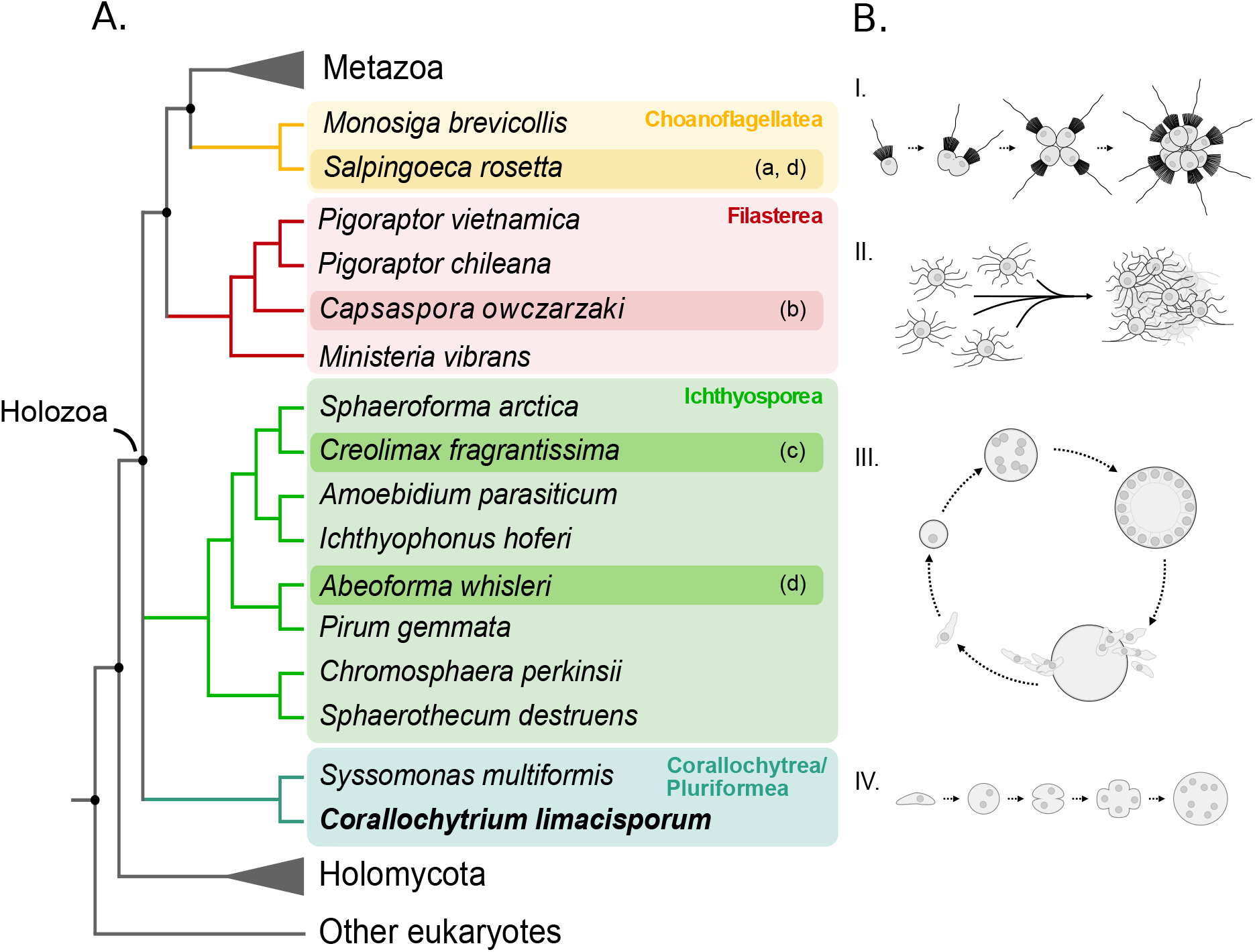
Unicellular holozoans and their phylogenetic position relative to Metazoa. **A**. A schematic phylogenetic tree of Holozoa based on (Torruella et al., 2015; Grau-Bové et al., 2017; Hehenberger et al., 2017). (a)-(d) Indicate the availability of transfection tools in these organisms. (a) (Booth et al., 2018); (b) (Parra-Acero et al., 2018) (c) (Suga and Ruiz-Trillo, 2013); (d) (Faktorová et al., 2020). B. Schematic representations of the developmental mode of representative species from each clade, emphasizing the multicellular stages. I. Clonal colony formation in *S. rosetta* (Fairclough et al., 2010; Dayel et al., 2011; Brunet and King, 2017). II. Aggregative multicellularity of *C. owczarzaki* (Sebé-Pedrós et al., 2013). III. Coenocytic growth of *C. fragrantissima* (Marshall et al., 2008; Suga and Ruiz-Trillo, 2013). IV. Putative coenocytic growth of *C. limacisporum* (Raghukumar S., 1987; Mendoza et al., 2002).

Comparative cell biology depends on the availability of genetic tools both to facilitate visualization of cellular structures and to perform perturbations. Key steps have been made recently in providing genetic tools for use in one choanoflagellate (Booth et al., 2018; Booth and King, 2020), two ichthyosporeans (Suga and Ruiz-Trillo, 2013; Faktorová et al., 2020) and one filasterean (Parra-Acero et al., 2018). However, the absence of such tools in Corallochytrea makes it currently impossible to undertake comparative cell biological studies across all holozoan clades. Given that Corallochytrea diverges substantially from the other holozoan lineages (Torruella et al., 2015; Hehenberger et al., 2017) it is now essential to develop genetic tools in this clade to allow comprehensive cell biological comparisons with broad taxon sampling.

The Corallochytrea currently includes only two known species: *Corallochytrium limacisporum* and *Syssomonas multiformis. C. limacisporum* was initially classified as a thraustochytrid fungi based on its morphology and the presence of some fungal-like molecular traits (Raghukumar S., 1987; Sumathi et al., 2006). Later on, however, it was shown to be an independent holozoan lineage, possibly a sister-group to the Ichthyosporea clade within the Holozoa (Torruella et al., 2015). Interestingly, it has some fungal as well as some animal features. For example, *C. limacisporum* is the only holozoan with chitin synthases homologous to the fungal and not to the animal genes (Torruella et al., 2015). Furthermore, *C. limacisporum* has numerous features that make it an attractive candidate for further functional analysis. First, it is the only corallochytrean with a completely sequenced and well-annotated genome (Torruella et al., 2015; Grau-Bové et al., 2017). Moreover, and in contrast to *S. multiformis*, it has the technical advantage of growing very fast and under axenic conditions in both liquid and agar medium, allowing easy screening and selection of individual transfected clones. These features together with its peculiar and under-studied biology make *C. limacisporum* a fascinating organism to develop as an experimentally tractable system.

The array of genetic tools and methodologies that can be established is diverse, but to analyze evolutionary cell biological questions, a robust transfection protocol together with an array of cell markers is essential. Protocols allowing stable transfection would be extremely valuable, as they allow transfected lines to be followed over long periods of time. Here, we describe a robust, stable transfection protocol in *C. limacisporum*, as well as a battery of cassettes for tagging different cellular components. Using these tools, we provide new insights into *C. limacisporum’s* life cycle including the characterization of some unexpected traits. We show that it exhibits two different paths for cell division—binary fission and coenocytic growth-demonstrating that the life cycle of *C. limacisporum* is more complex than previously thought (Raghukumar S., 1987). Interestingly, we also report two features of *C. limacisporum*, which are considered rare in holozoans and eukaryotes (Martin et al., 2008; Alberts et al., 2002; Gladfelter, 2006). First, the decoupling of cytokinesis and karyokinesis in binary fission, and second, the occurrence of asynchronous nuclear divisions during coenocytic growth. The possibility to expand functional studies of these features in *C. limacisporum* will undoubtedly contribute not only to a better characterization of this independent holozoan group, but also to understanding both the evolution of metazoan cell biological features and the emergence of metazoans.

## RESULTS

### A robust protocol for stably transfecting Corallochytrium limacisporum

The first step towards establishing stable transfection is to develop a selection system. Therefore, to assess antibiotic sensitivity in *C. limacisporum*, we assayed a diverse panel of drugs, including commonly used antibiotics, antifungals and herbicides (Supp. Table 1). We observed that *C. limacisporum* is susceptible to puromycin, which efficiently killed the cells after three days at a concentration of 300 μg/mL. Based on this finding, we cloned the corresponding resistance gene, puromycin N-acetyltransferase (*pac*), which we fused with the gene encoding the mCherry fluorescent protein to provide a double selection system for positive cells: drug resistance and fluorescence. The actin and tubulin promoters are known to be constitutively active in other systems (Angelichio et al., 1991; Joung and Kamo, 2006; Hernandez-Garcia and Finer, 2014). However, to date, no regulatory sequences have been described in *C. limacisporum*. Therefore, to increase the chances of obtaining a functional gene-regulatory sequence, we targeted both of these as putative promoter regions in two recombinant plasmids (Fig. 2A): CAMP (***C**orallochytrium* **A**ctin **M**cherry **P**ac) and CTMP (***C**orallochytrium* **T**ubulin **M**cherry **P**ac).

**Figure 2.**
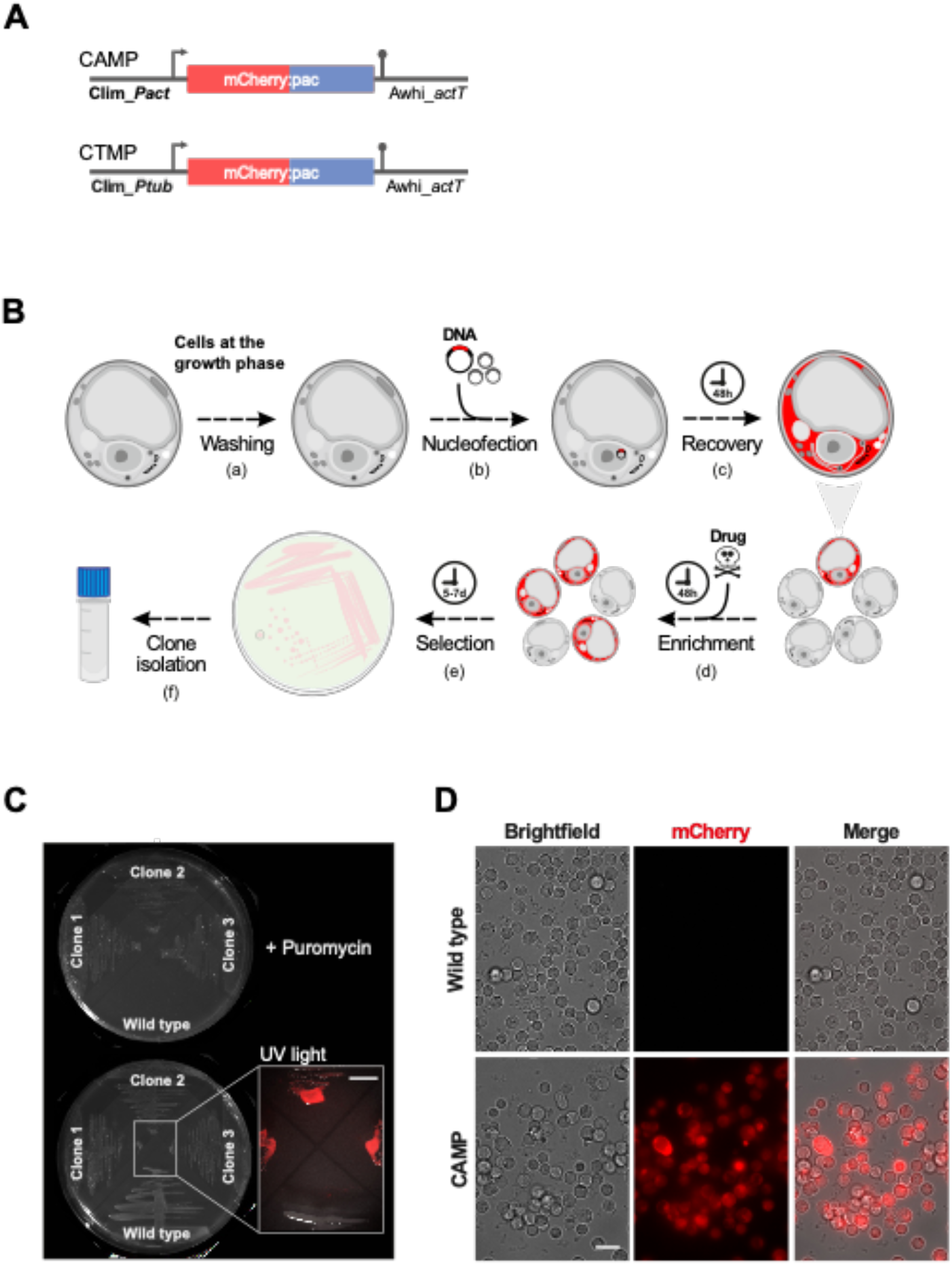
Transfection of *Corallochytrium limacisporum* by nucleofection. **A.** Transfection constructs: mCherry fused to puromycin resistance gene (*pac*) under either promoter variant: actin (p_act) – CAMP - or tubulin (p_tub) - CTMP. **B.** Steps of the protocol: (a) cells at the growing phase (1.5×10^6^ cells) are washed with 1XPBS to remove culture medium; (b) cells are mixed with transfection construct (10 μg) and carrier DNA (40 μg) and 20 μl of Lonza P3 commercial buffer and subjected to transfection program in Lonza 4D-Nucleofector™ System (EN-138), (c) cells are then transferred to 1 mL of culture medium for recovery period of 48h; (d) puromycin at the concentration of 300 μg/mL is added for selection and enrichment of positive cells; (e) after 48h cells are transferred on puromycin-rich marine agar plates and grown for 5 - 7 days; (f) pink-rounded colonies can be isolated for further manipulations (see *Material and methods* for more details). **C. & D.** Examples of established clonal lines of *C. limacisporum India* transfected with CAMP plasmid and grown in the presence of puromycin either on solid medium (**C**) or in liquid medium (**D**). Wild type (not transfected) lines do not grow in the selective medium neither they show fluorescence in the non-selective medium.

To deliver recombinant DNA into *C. limacisporum* cells, we initially tested various chemical and physical transfection protocols. In this initial screen, we obtained a handful of transfected cells when using electroporation-based methods. We first used the Neon electroporation system (Invitrogen) (protocols.io: dx.doi.org/10.17504/protocols.io.hmwb47e), which allows high voltage and multiple pulses to be applied, and which had been previously successful in our hands for transfection of the ichthyosporean *Creolimax fragrantissima* (Suga and Ruiz-Trillo, 2013)(Fig. 1). Nevertheless, our attempts to further optimize this protocol did not result in either higher efficiency or reproducibility, leading us to focus our efforts instead on another electroporation-based system, 4D-nucleofection (Lonza^®^), which had been applied successfully in the choanoflagellate *S. rosetta* (Booth et al., 2018) and the ichthyosporean *A. whisleri* (Faktorová et al., 2020) (Fig. 1).

We used the CAMP plasmid (Fig. 2A) to optimize the 4D-nucleofection protocol (Fig. 2B and protocols.io: dx.doi.org/10.17504/protocols.io.r5ud86w). Based on analysis of cell viability, we chose an optimal density for transfection of 1.5×10^6 cells/reaction at their exponential growth phase (2-day-old culture) (Fig. S1A). We additionally improved the viability of the cells by washing the medium away with 1XPBS before applying the electric pulse. Results showed that 10 μg/mL of the recombinant plasmid DNA provided the largest number of positive cells (Fig. S2A), which was significantly enhanced by the addition of 40 μg/mL of plasmid carrier DNA (pUC19) (see efficiency analysis below). We then tested the different electric parameters combined with the different transfection buffers provided by Lonza^®^ and we observed that the combination of buffer P3 and code EN-138 proved to be the best for successful transfection of *C. limacisporum*. Finally, we also observed that adding the medium immediately after transfection was crucial for maximum cell recovery. We further extended the recovery period for 48 h, after which we applied puromycin selection. Selection was performed either by plating the recovered cells on marine agar for clonal isolation or by successive passages of the transfected cells in liquid medium. We show different examples of established clonal lines with the described transfection protocol (Fig. 2B) and the CAMP plasmid (Fig. 2A). As mentioned above, the CAMP plasmid employs a double selection system for positive cells: by drug resistance, growing the given culture in the presence of puromycin, and by mCherry fluorescence. Transfected cells can be selected either clonally in marine agar plates (Fig. 2C) or in liquid medium (Fig. 2D). Independent transformed lines can be preserved following the cryopreservation method that we optimized for *C. limacisporum* (see Material and Methods).

To determine transfection efficiency, we quantified the number of positive cells for each of the engineered constructs, CAMP and CTMP, using flow cytometry. We used untransfected wild type *C. limacisporum* cells to define the gating for the negative population, and all events above this threshold were considered as a positive population (Fig. S2B). We performed the transfection experiments with and without the addition of a carrier DNA (pUC19). We quantified the efficiency of transfection 48h post-transfection for the two available strains of *C. limacisporum, India* and *Hawaii* (Fig. S2C), which molecular data indicates are almost identical (Torruella et al., 2015). Based on six replicates from two independent experiments, we obtained an average efficiency for *C. limacisporum India* of 0.15%± 0.05 (mean±s.d.) and 0.22%±0.5 for CAMP and 0.19±0.13 and 0.3%±0.08 for CTMP, without and with carrier DNA, respectively. *C. limacisporum Hawaii* efficiency was 0.1±0.08 and 0.16±0.08 for CAMP and 0.2±0.05 and 0.34±0.14 for CTMP, without and with carrier DNA, respectively (Fig. S2C). As the above numbers indicate, the percentage of transfected cells obtained for both *C. limacisporum* strains were in the same range. Therefore, the rest of the reported experiments were done in either strain (indicated in figure legends). Since the addition of carrier DNA (pUC19) increased the number of transfected cells, we performed all the transfection experiments with the addition of pUC19. Moreover, we obtained a slightly higher percentage of positive cells when transfecting with the CTMP construct. To better understand this difference, we examined the strength of both promoters (actin and tubulin) (Fig. S2D). In particular, we measured the mean intensity of the expression of the mCherry protein under each promoter, and the obtained values were normalized to the plasmid copy number in each strain, as measured by qPCR (see Material and Methods for details). The results showed that the tubulin promoter is three times stronger than the actin promoter, which confirmed our microscopic observations. These results were taken into consideration when designing the additional constructs reported here. Taken together, our results show that reliable, stable transfection of *C. limacisporum* can be achieved using these protocols.

### Live imaging of C. limacisporum by labelling cellular components with fluorescent markers

The first step in comparative cell biology is to visualize and localize the cellular structures of interest in a given system. Therefore, we generated three constructs to enable live imaging of the plasma membrane, actin cytoskeleton and nucleus of *C. limacisporum* in addition to CAMP and CTMP, which label the cytoplasm (Fig. 3). Imaging of CAMP- and CTMP-transfected lines allowed the first more detailed observations of cellular morphology in *C. limacisporum*. Cytoplasmic expression of mCherry showed that the cytoplasm is restricted to the cell periphery, generating a ‘crescent-like’ shape in the majority of cells (Fig. 3A and Fig. S3A_I). This pattern suggests the presence of a large vacuole, occupying on average 65% of the cell’s volume. We confirmed the presence of this vacuole using pulse-chase experiments with the lipophilic plasma membrane dye FM4-64 (Fig. S3B). Interestingly, in some cases, the cytoplasm is present also on the other side of the vacuole, resulting in the vacuole occupying only 40% of the total cell volume (Fig. S3A_II). In these cases, it seems most likely that the cell is preparing to divide (Fig. S3A_III).

**Figure 3.**
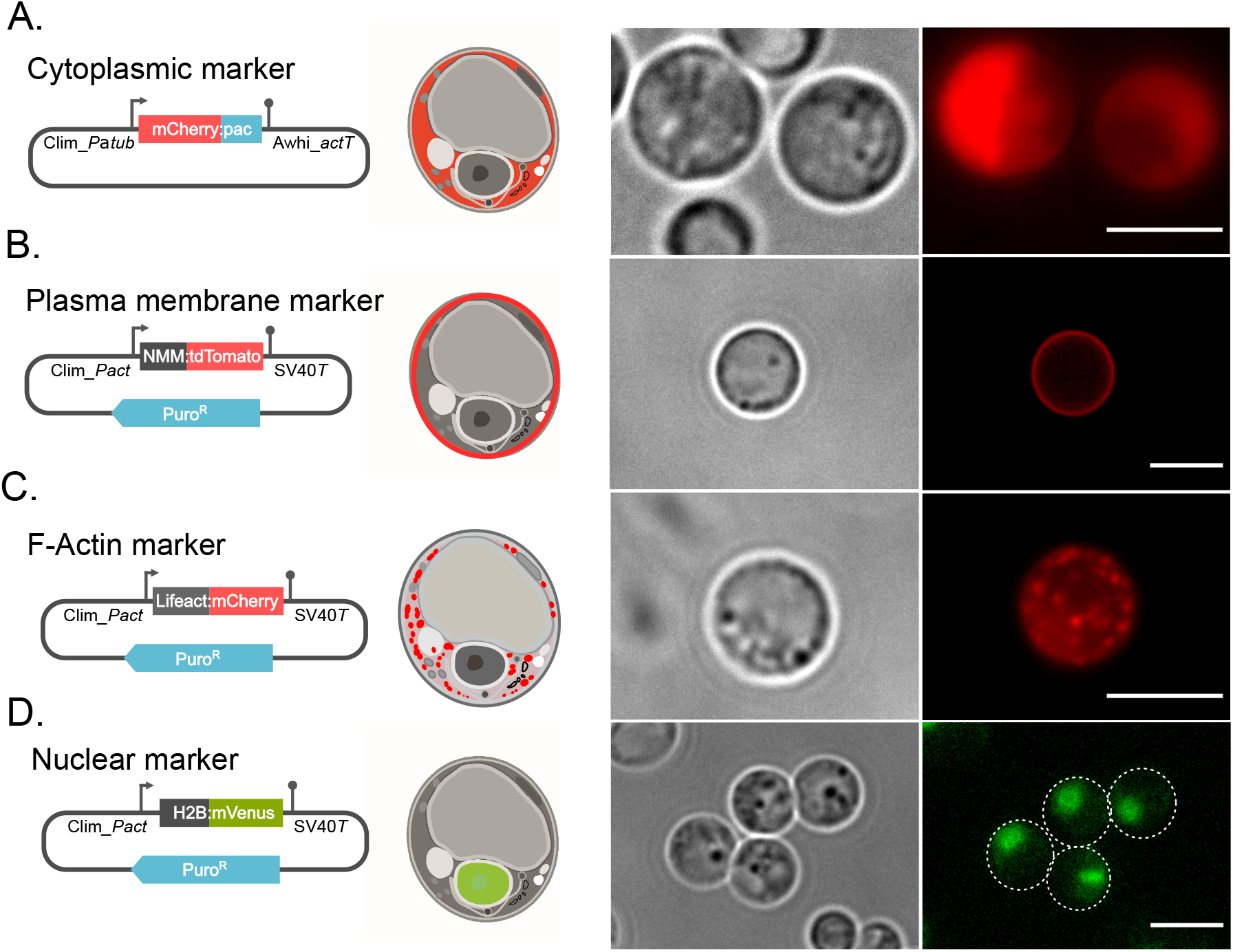
Expression cassettes for subcellular tagging and live imaging of transfected *Corallochytrium* cells. Live imaging of cells transfected with different cellular markers tagging key cellular components. Images are complemented by drawings of expression cassettes (left) and schematic interpretations of marker localization (middle). **A.** Cytoplasmic marker cassette expressing mCherry fluorescent protein. **B.** Plasma membrane marker cassette expressing *C. limacisporum* Src NMM fused to mCherry. **C.** F-Actin marker cassette expressing LifeAct fused to mCherry. **D.** Nuclear marker cassette expressing *C. limacisporum* histone H2B fused to mVenus. All cassettes contain the puromycin resistant gene (*pac*). Cells were imaged using wide-field fluorescence microscopy and pseudo - colored with ImageJ software. Scale bars: 5μm.

To label the plasma membrane and visualize its remodelling during key cellular processes such as cell division, we generated a construct (pClimDC-NMM:tdT) expressing an endogenous membrane-binding motif fused to the fluorescent protein tdTomato. We chose the predicted N-myristoylation motif (NMM) from the *Src* tyrosine kinase orthologue (Gene ID Clim_evm93s153), which is commonly used to direct fluorescent protein markers to the eukaryotic plasma membrane and which has been previously used successfully in *Capsaspora owczarzaki* (Parra-Acero et al., 2018). We were able to visualize the plasma membrane of live *C. limacisporum* cells (Fig. 3B) and observe its dynamics during cell division (Movie 1). This tool allowed us to precisely identify when cytokinesis was completed (see below).

In order to label the actin filaments of the cytoskeleton, we designed a construct (pClimDC-LifeAct:mCh) for the expression of mCherry fused to LifeAct, a 17-amino-acid peptide that binds specifically to filamentous actin (Riedl et al., 2008). With this construct, we were able to clearly visualize actin organization. Actin filaments appeared as patches that were evenly distributed throughout the cell (Fig. 3C). Because this was the first time that actin organization had been visualized in *C. limacisporum*, we validated these results by phalloidin staining in fixed cells (Fig. S4), which confirmed the pattern obtained with the F-actin marker. This construct will be useful for future experimental designs to study cytoskeleton dynamics during various cellular processes.

Finally, we generated a construct (pClimDC-H2B:mV) to label the nucleus by tagging the endogenous histone H2B (Gene ID Clim_evm20s1) with the mVenus fluorescent protein. Lines transformed with this construct showed mVenus signal specifically localized to the nucleus (Fig. 3D). Nuclei were well defined and in general displaced from the centre, likely due to the presence of the central vacuole (see above and Fig. 3D). Visualization of the nuclei in asynchronous cultures of *C. limacisporum* revealed different nuclear configurations corresponding to different life stages (see next section).

All of the above constructs were used to generate the stable lines that are shown in Fig. 3 (always from single isolated colonies). We verified the genomic integration of the constructs pClimDC-LifeAct:mCh and pClimDC-H2B:mV by PCR (Fig. S5). Importantly, none of the constructs displayed toxicity, and the corresponding stable lines were successfully propagated for several generations. Overall, our data show that these constructs can be used to specifically label cellular components in *C. limacisporum*.

### Live observation of lines carrying a nuclear marker allows reconstruction of the life cycle of C. limacisporum

Our current understanding of the life stages of *C. limacisporum* has been limited to observation of live cultures by bright field microscopy or of fixed cells (Raghukumar S., 1987; Torruella et al., 2015). However, these approaches have proved insufficient to reconstruct the complete life cycle of *C. limacisporum* and the specific characteristics of its different stages. For example, previous observations suggested the existence of a multinucleate stage that would resemble the coenocyte in Ichthyosporea, but without a nuclear labelling this had not been demonstrated. Therefore, to identify the number of nuclei in each of the stages of *C. limacisporum*, we imaged lines transformed with the pClimDC-H2B:mV plasmid (Fig. 3D). We discerned cells with one, two, three, four, eight or more nuclei in the same cell (Fig. 4A). Thus, the life cycle of *C. limacisporum* includes a coenocytic stage.

**Figure 4.**
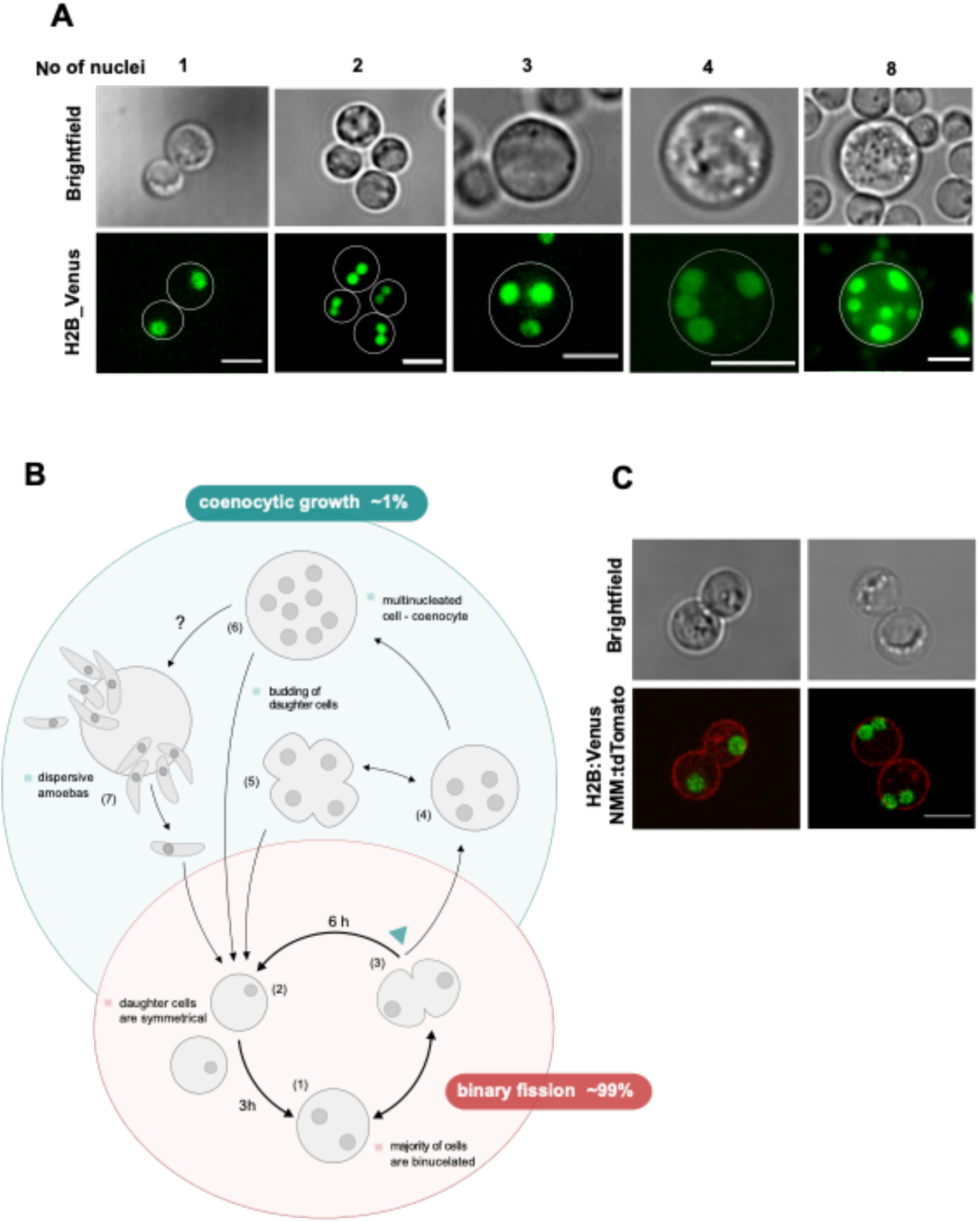
A comprehensive understanding of *C. limacisporum* life cycle. **A.** Live imaging of *C. limacisporum Hawaii* nuclei (here 1, 2, 3, 4, and 8 nuclei) confirming the presence of coenocytic growth in this organism. Cells with 2, 3 and 8 nuclei were imaged with wide-field fluorescent microscopy and cells with 1 and 4 nuclei with Spinning disc confocal microscopy. All images are presented as the maximum projection of Z-stacks, processed with ImageJ. Scale bars: 5μm. **B.** Scheme of the proposed life cycle of *C. limacisporum*. Reproduction in *C. limacisporum* occurs mainly through binary fission, during which a binucleated cell (1) divides into two, symmetrical, uninucleate cells (2). This process occurs in 99% of cases (red background). Binucleate cell can form two lobes (3) that can lead to cellular division (forming two monoucleate cells), or it can reverse towards spherical cells and continue dividing its nuclei further forming quadrinucleated cell (4). Quadrinucleated cell can often form a quatrefoil-like shape (5) (similar to bilobed cell), generating either four mononucleate cells or return to spherical shape and divide to eight (6), twelve and up to thirty-two nuclei coenocyte. Dispersive amoebas can be released from a coenocyte (7). The coenocytic growth occurs in about 1% of the cases (blue background). **C.** Representation of *C. limacisporum* cells before and after the completion of cytokinesis. Pictures on the left show cells before the completion of the cell division. Picture on the right shows two cells after the cytokinesis was completed. This distinction was important for differentiating between bilobed cells and (Fig.4B (3)) two separated daughter cells that are attached one to another (Fig.4B (2)→(1)). Images are presented as the maximum signal projection of Z-stacks and were acquired with confocal microscopy. Scale bar: 5μm.

Next, we reconstructed the life cycle of *C. limacisporum* by live imaging and time-lapse movies (scheme of *C. limacisporum* life cycle: Fig. 4B). Interestingly, the most commonly observed cells are binucleate (Fig. 4A_2, Fig. 4B (1)). Both nuclei are of similar size and shape, and they are localized next to each other (Fig. 4A_2). Live imaging and time-lapse experiments revealed that binucleate cells can have two fates: either dividing into two mononucleate cells (Fig. 4B (2)), Movie 1) via binary fission (99% of cases), or less frequently, increasing in nuclei number and cell size to form a coenocyte (Fig. 4A_8, Fig. 4B (6), Movie 2 - arrow 1, Fig. 4_8). During binary fission, when one cell gives rise to two identical daughter cells, we observed some intermediate stages. Before dividing, the binucleate cell forms two lobes (Fig. 4B (3), Fig. 4C, Movie 3 / time 10:35 - arrow 4). This bilobed cell can proceed towards either a round of cell division, with the subsequent birth of two symmetrical daughter cells (Fig. S6A), or return to a spherical binucleate cell that will most likely go through another round of nuclear division, giving rise to a cell containing four nuclei (Fig. 4B (4), Fig. 4A_4 and Movie 2 / time 16:10 - arrow 2). These cells can form quatrefoil-like cell (Fig.4B (5), Movie 2 / time 22:55 - arrow 3) that, similar to bilobed cells, can either undergo cytokinesis to complete cell division or return to rounded cells and continue multiplying their nuclei to eight (Fig. 4B (6), Fig. 4A_8). Cells with more than four nuclei that continue dividing will give rise to coenocytes, which in all observed cases, gave rise to mononucleate cells in a budding-like manner (Movie 3/ 13:10 - 15:40 – arrow 5) or releasing swimming limax-shaped amoebas (Fig. 4B (7)), Movie 4). Even though coenocytic growth is significantly less frequent than binary fission, it is not uncommon to encounter coenocytes in the culture. This can be explained by a difference in the cell size of coenocytes vs non-coenocytic cells at the regular density of the culture. We quantified that a cell with four nuclei is on average 6.5 μm in diameter, which is 20 – 30 % larger than non-coenocytic cells (bi- or mononucleate) (Fig. S6B). The difference in cell size between cells (mononucleate, binucleate and quadrinucleate) can be appreciated in Movie 2 / time 00:00. The size of a non-coenocytic cell is around 4.5 μm, increasing up to 15 - 20% before dividing (Fig. S6C). Interestingly, we also observed coenocytic cells with odd numbers of nuclei. In particular, cells with three (Fig. S7A) or seven nuclei (Fig. S7B). These results indicate that nuclear division can occur asynchronously, which should be further investigated. Overall, establishing transformed lines with labelled nuclei has been key to reconstructing the life cycle of *C. limacisporum*, as well as discovering some unusual traits, such as a binucleate stage, the existence of a coenocyte and the potential to divide nuclei asynchronously.

### Time of nuclear and cellular division in C. limacisporum

In actively dividing animal cells, nuclear division (karyokinesis) is usually closely followed by the separation of cell cytoplasm (cytokinesis) to produce two daughter cells. Our observation that *C. limacisporum* cells are mainly binucleate indicates that cell division does not occur immediately after nuclear division in this species. To further understand the time difference between karyokinesis and cytokinesis, we analysed lines that were co-transfected with pClimDC-H2B:mV and pClimDC-NMM:tdT and to allow simultaneous visualization of the nucleus and plasma membrane (Fig. 5A and Movie 1). We considered the start point of the cycle when a binucleate cell gave rise to two symmetrical, mononucleate daughter cells. These mononucleate cells would soon divide their nuclei once again to obtain the phenotype depicted in Fig. 5A. Based on 60 measurements, we calculated that nuclear division takes on average 2 hours, whereas it takes up to 6 hours for cell division to be completed (Fig. 5B). These findings revealed that the binucleate stage lasts approximately 4 hours and suggested that, unlike in most eukaryotic cells, nuclear division is decoupled from cytokinesis in the binary fission of *C. limacisporum* (Fig. 5B). Additionally, our data showed that binucleate cells are larger, on average, than mononucleate cells (Fig. S6B). Measuring the variation in cell size in relation to the number of nuclei revealed that the larger (binucleate) cell stage occupies about 2/3 of the duration of the cycle, whereas smaller (mononucleate) cells account for the remaining 1/3 of the binary fission cycle (Fig. 5C). These data can serve as a starting point towards an understanding of the machinery responsible for this unusual cell cycle in *C. limacisporum*.

**Figure 5.**
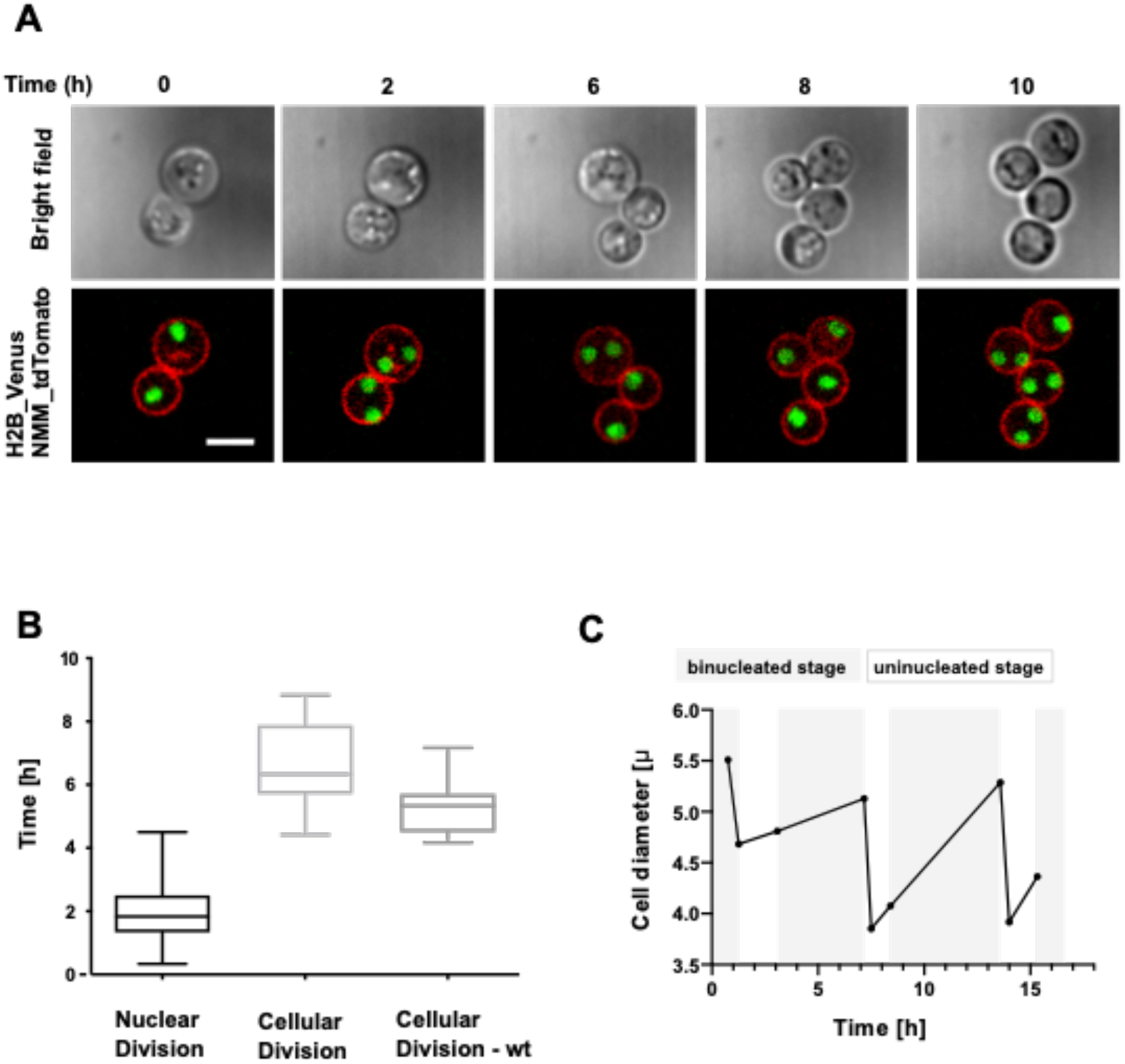
Decoupling of nuclear and cellular division in *C. limacisporum*. **A.** Images of *C. limacisporum Hawaii* transformants with labelled nuclei (green) and plasma membrane (red). Images were taken every 5 minutes for 23h using spinning disc confocal microscopy. Complete movie can be seen in Supplementary Movie 1. Scale bar: 5μm. **B.** Analysis of nuclear and cellular division time in *C. limacisporum Hawaii*. Average nuclear division time equals 2h (number of nuclear divisions measured: 64), whereas cellular division time equals 6h (number of cellular divisions measured: 46), indicating that cytokinesis is decoupled from karyokinesis. **C.** Representation of time range in which cells are binucleated (gray) or mononucleate (white) against their respective cell size. Cell diameter was calculated using ImageJ software.

## DISCUSSION

### Holozoa emerges as a unique clade to analyze cell biological questions

Our understanding of the evolution of cellular processes in eukaryotes has, until recently, been limited by the lack of established experimental systems in most parts of the eukaryotic tree of life. For many years, we only had genetic tools in some fungi and animals. More recently, tools were also developed in some protists, most of them of clinical relevance (Lander et al., 2016). However, the reduction of sequencing costs and the advent of molecular tools, has allowed a diversification of efforts to develop tools in more eukaryotic taxa (Faktorová et al., 2020).

To perform useful comparative cell biological analyses, ideally, genetic tools should be available for different taxa from the same clade. One should look for biologically relevant taxa situated in a key phylogenetic position in the tree of life. For example, to understand the evolution of cell biological features in animals or fungi, one should aim to have experimentally tractable organisms in different fungi and animal groups, as well as in their various unicellular relatives.

The case of animals is unique, given that their cell biology is relatively well known but has not yet been compared with the different unicellular taxa with which they share a common ancestor. Indeed, several unicellular lineages have been shown to be closely related to animals, each one with its own cell morphology and developmental characteristics. Thus, to analyze the evolution of cell features from protists to animals in the broadest possible way, one should be able to investigate organisms from each of the different unicellular linages closely related to animals. In this study, we have successfully established an experimentally tractable system in the Corallochytrea. The availability of a reliable method for stable transfection of *C. limacisporum* will now make it possible to perform taxon-rich comparative experimental analyses among all described holozoan lineages, from animals to choanoflagellates, ichthyosporeans, filastereans and, now, corallochytreans.

### C. limacisporum has a complex life cycle consisting of both binary fission and coenocytic growth

The first investigation of *C. limacisporum* was done in 1987 by Raghukumar (Raghukumar S., 1987). In that study, the linear life cycle was proposed to be initiated by a limax-shaped amoeba stage that eventually becomes a mononucleate cell. This cell then undergoes successive rounds of binary cell division, after which cells remain attached to each other, forming up to a 32-cell stage. However, our own microscopy observations have now shown that the life cycle of this taxa may be more complex.

Understanding the full breadth of life cycles among Holozoa is of clear evolutionary interest, given that some of those life cycles may have been key to the origin of animal cell types or even the origin of animals. It is also relevant from an ecological and economic point of view, since some of the holozoans have complex life cycles that include a parasitic stage in animals of economic relevance, such as fish and amphibians (Rowley et al., 2013).Thus, we decided to take advantage of the tools developed here to obtain a better understanding of the life cycle of *C. limacisporum*.

Our stably transfected lines allowed us to discern the mononucleate, binucleate and coenocytic life stages in *C. limacisporum* and follow their dynamics throughout the life cycle. Our results demonstrate that the life cycle of *C. limacisporum* is non-linear and is composed of two different paths. Firstly, there is a well-coordinated binucleate stage reached by binary fission, where a binucleate cell gives birth to two symmetrical mononucleate daughter cells (99% of cases in our culture conditions). Secondly, *C. limacisporum* exhibits coenocytic growth, where a binucleate cell divides its nuclei without accompanying cytokinesis (1% of cases in our culture conditions), with a possibility of reaching up to 32 nuclei (Raghukumar S., 1987). Interestingly, those developmental pathways are also present in other unicellular Holozoa. Binary fission, the common mode of division in most holozoans, resembles the one occurring in choanoflagellates and filastereans. In contrast, coenocytic growth, considered as a simple multicellular stage, is a characteristic of ichthyosporeans. However, to our knowledge, this is the first example of a holozoan in which the two modes of development are both present. Unlike the process reported in ichthyosporeans, coenocytic growth in *C. limacisporum* is a facultative developmental mode that occurs rarely, at least in our culture conditions, and seems to be triggered either by external or internal factors. Which factors govern the decision, and whether they are genetic, environmental or both, is unclear and will require further investigation.

### Nuclear and cell division are decoupled in C. limacisporum

Binucleation is an uncommon trait amongst eukaryotes. Rare instances are found in the metamonad *Giardia lamblia* (Bernander et al., 2001), the thraustochytrid fungi of the *Aurantiochytrium* species (Ganuza et al., 2019) and in some microsporidians (Keeling and Fast, 2002). Two nuclei (the micro and the macro nucleus (Prescott, 1994), are also a characteristic feature of some ciliates. Similar to *G. intestinalis*, and in contrast to ciliates, the two nuclei observed in *C. limacisporum* are of similar size and, presumably, have the same functions. However, while *G. lamblia* alternates between binucleate trophozite and quadrinucleate cystic forms (Bernander et al., 2001), our data show that *C. limascisporum* goes through a mononucleate stage, since it eventually completes the cell cycle by performing cytokinesis through a decoupled mechanism. Furthermore, in contrast to *Aurantiochytrium* species and *G. lambia, C. limacisporum* only reaches a quadrinucleate cell when following coenocytic growth and not during binary fission. Moreover, *C. limacisporum* does not show the wide variety of forms in its life cycle seen in *Aurantiochytrium* species (Ganuza et al., 2019). The distinctive traits suggest that the binucleation observed in *C. limacisporum*, due to the decoupling of nuclear and cell division, is a distinct process worthy of further exploration. It is tempting to speculate that the delay in cytokinesis and the bi-nucleated stage are related to either environmental sensing or to a specific timing for molecular adjustments of the genomic content of each nucleus. Further studies into the dynamics of the nuclear envelope and the microtubule machinery are required to better understand the mechanisms involved, and more in-depth analysis of the culture behavior will be required to fully elucidate the nature of this unusual binary fission.

Cell growth is determined by the relationship between increasing cell size and the frequency of cell division. Coordination of cell division through mechanisms that sense cell size occurs in a range of taxa (Turner et al., 2012; Tzur et al., 2009), and it has recently been found that the cell size depends on nutrient concentration in the ichthyosporean *Sphaeroforma arctica* (Ondracka et al., 2018). In our culture conditions, the cell volume in *C. limacisporum* increases by 20-30% before division. However, we did not identify a required cell size to trigger cell division. We have taken into account that the cell environment in our cultures changes over time with the reduction in nutrients and increased crowding of the culture. Therefore, these measurements were performed in the initial states of the culture when nutrients are not limiting and the culture is at optimal density. Our results indicate, therefore, that there is no size-dependent mechanism controlling cell division. Nevertheless, we found that quadrinucleate cells are, on average, twice as large as mononucleate cells, indicating that a correlation between cell size and number of nuclei does exist.

### C. limacisporum coenocytes can divide asynchronously

Our observations by live imaging demonstrate that multinucleate cells in *C. limacisporum* are formed by mitotic nuclear division and not by cell fusion. Nuclear division is usually coordinated within a common cytoplasm (Rao and Johnson, 1970), meaning that synchronous division of nuclei would normally be expected within a given cell. This is the case in other unicellular holozoans, such as *S. arctica* (Dudin et al., 2019) and *C. fragrantissima* (Suga and Ruiz-Trillo, 2013). In contrast, asynchronous cell division is a rare phenomenon, so far only described in a few eukaryotic taxa, such as the filamentous fungi *Ashbya gossypi* (Gladfelter, 2006). Nuclei in *A. gossypi* can be in different cell cycle stages despite their close physical proximity. The molecular mechanism proposed to be behind this asynchrony is based more on the action of inhibitors of cyclin-dependent kinases than on controlling the amount of cyclin protein (Gladfelter, 2006). Alternative mechanisms for independent nuclear regulation have been described in response to other phenomena, such as DNA damage and the presence of unattached chromosomes during mitosis (Rieder et al., 1997; Demeter et al., 2000;). However, the study of how asynchronous nuclear division is regulated in multinucleate cells is limited to relatively few systems. Our data shows that nuclear division in *C. limacisporum* coenocytes can either happen synchronously or asynchronously, serving therefore as a good model to understand the regulation of both types of nuclear division.

## CONCLUSIONS

Here, we report the successful development of a new experimentally tractable organism among unicellular holozoans, the corallochytrean *C. limacisporum*. We have established a protocol for stable transfection together with genetic constructs for live imaging. Using these tools, we have reconstructed the life cycle and described biological features that seem to be specific to *C. limacisporum* within the Holozoa. Together with experimental organisms from Choanoflagellatea (Booth et al., 2018), Filasterea (Parra-Acero et al., 2018) and Ichthyosporea (Suga and Ruiz-Trillo, 2013) the establishment of genetic tools in *C. limacisporum* completes a functional platform to address questions related to the evolution of cellular mechanisms and the development of multicellularity. Importantly, *C. limacisporum* provides the opportunity to further investigate the factors behind different developmental routes (binary fission or coenocytic growth), as well as offering a promising model to study the mechanisms behind decoupling of karyokinesis from cytokinesis and the basis of asynchronous nuclear division.

## ACKNOWLEDGMENTS

We thank the UPF Flow Cytometry Core Facility for their constant kind assistance. We thank the CRG Advanced Light Microscopy Unit for their help in acquisition of confocal movies and images. We thank David Booth for his advice on the transfection protocol and generous sharing of carrier plasmid DNA. We thank Meritxell Antó for technical support. We thank Omaya Dudin and Núria Ros i Rocher for their support in analyzing microscopy images and the MulticellGenome laboratory for fruitful discussions. This work was funded by the Gordon and Betty Moore Foundation’s Marine Microbiology Initiative for establishing Emerging Model Systems Grant Number: 4973.01.

## AUTHOR CONTRIBUTION

AK, SN, and EC designed the study and set up the methodology. AK, SN and CA designed and built the DNA constructs. AK, SN performed the transfection protocol development and transfection, selection and microscopy experiments. EO performed the bioinformatic analysis of transfected lines. I.R.-T. and EC. provided supervision. AK and EC wrote the original draft. All authors reviewed and edited the manuscript.

## MATERIALS AND METHODS

### Cell culture and growth conditions

*Corallochytrium limacisporum* strains ‘*Hawaii*’ (isolated in Kāne’ohe Bay, Hawaii (Torruella et al., 2015) and ‘*India*’ isolated in Kavaratti Island, India (Raghukumar S., 1987) were grown axenically in 25 cm^2^ flasks (Falcon^®^ VWR, #734-0044) with 5 ml of marine broth (Difco™, # 2216), in an air incubator at 23 ºC. Cultures were transferred to fresh MB medium approximately once a week and were maintained in the lab conditions for months before the analysis were completed. For mid-term storage, and / or for clonal selection cells were grown on solid marine agar (Difco™, #2216), in 5 cm diameter plastic Petri dishes with or without the addition of 300 μg/mL of puromycin (Sigma Aldrich #P7255), as indicated in the text.

### Determination of growth curves

Cells from one-day-old culture - at the mid-exponential growth phase - at an initial cell density of 1×10^5^ cells / ml were seeded in 12-well plates (Nunc/DD Biolab, #55428) with 1 ml marine broth and incubated at 23 ºC. At each time point, triplicate cultures were scraped and counted using a Neubauer chamber. The calculated numbers of cells/ml were recorded in Excel and used to plot the growth curves (Fig. S1A).

### Sensitivity and resistance to antibiotics

Wild-type *C. limacisporum* cultures at an initial density of 2 x10^6^ were exposed to a panel of different antibiotics (at concentrations ranging from the solubility limit of each drug). The same volume of solvent, corresponding to the maximum antibiotic concentration, was added to cells as a control. Cells were monitored for over a week and assessed by microscopy. Concentrations tested were 50 and 100 μg/ml in the indicated solvent by manufacturer unless indicated otherwise, such as: puromycin 100 – 500 μg/ml, benomyl 20 - 500 μg/ml, carboxine 20 - 300 μg/ml, nourseothricin 10-50 μg/ml and fluoretic acid 250 - 30 μg/ml (Fig. S1). After determining that *C. limacisporum* cells were susceptible to puromycin, we tested a range of concentrations, from 100 μg/ml - 500 μg/ml. The concentration of puromycin that visibly affected the culture in the shortest amount of time was 300 μg/ml. Hence, this concentration was chosen for selection of resistant cells.

### Construction of expression vectors

*C. limacisporum* transfection vectors were built using pCR2.1 plasmid (Invitrogen) as the backbone. First, pCR2.1 was reduced in size by taking out ~1.5kb of the original vector, containing the f1 origin of replication and part of the kanamycin resistance cassette. This was performed by digesting the circularized plasmid with *ApaI* and *NcoI*, followed by a fill-in reaction using DNA polymerase I large (Klenow) fragment (New England Biolabs), and re-ligation with T4 DNA ligase (New England Biolabs). This procedure give rise to pCR2.1Δf1/Kn^R^, of 2377 bp.

Genomic DNA from *C. limacisporum* and *Abeoforma whisleri* were extracted as in Grau-Bové et al., 2017 (Grau-Bové et al., 2017) and used as a template for endogenous elements of transfection vectors. *C. limacisporum* transfection vectors bear either the putative promoter region of endogenous actin (Clim_evm35s218) or α-tubulin (Clim_evm7s11) genes, and the terminator (3’UTR) region of the *A. whisleri* actin gene (Awhi_g50), driving the constitutive expression of the fluorescent protein mCherry fused to the puromycin N-acetyltransferase gene (*pac*). The 3’UTR region from *A. whisleri* actin gene (352 bp) was amplified by PCR using primers 1 and 2 (see Table S9 for primer sequences), and cloned into pCR2.1Δf1/Kn^R^ using the *XhoI* and *XbaI* restriction sites. Later, the *pac* gene was amplified from pBABE-puro plasmid (kindly provided by Jay Morgenstern and Hartmut Land, Addgene plasmid #1764) (Morgenstern and Land, 1990) using primers 3 and 4, and cloned between the *NotI* and *Xho*I restriction sites. The sequence coding for the fluorescent protein mCherry was amplified from pONSY-mCherry plasmid (Parra-Acero et al., 2018) using primers 5 and 6, and cloned in-frame, upstream of *pac* gene using *EcoRV* and *Not*I restriction sites. Finally, the upstream sequences of *C. limacisporum* actin (Clim_evm35s218 - 551 bp) or α-tubulin (Clim_evm7s11 - 764 bp) genes were amplified using primers 7 and 8 or 9 and 10, respectively, and inserted between *HindIII* and *EcoRV* sites. The final constructs were named CAMP (Addgene #104446) and CTMP (Addgene #129560), respectively (Fig. 2A). Both plasmids allow to track transfected cells by fluorescence microscopy, while providing resistance to puromycin.

For the construction of the double-cassette vector, we first replaced the 3’UTR region of *A. whisleri* actin gene in CTMP by the simian virus 40 (SV40) terminator, containing the SV40 polyadenylation signal, to avoid any putative side effect caused by *A. whisleri* 3’UTR (discussed in *Ocaña et al in preparation*). For this purpose, the SV40 terminator sequence was PCR-amplified from plasmid pSBtet-GP (pSBtet-GP was a gift from Eric Kowarz, Addgene plasmid # 60495) (Kowarz et al., 2015) using primers 11 and 12, and cloned into CTMP between the *XhoI* and *XbaI* restriction sites. This cloning give rise to CTMP-SV40T plasmid, which probed to be fully functional as the original version (data not shown). Next, the *pac* gene was amplified by PCR from pBABE plasmid using primers 13 and 4, and cloned between the *XmaI* and *XhoI* restriction sites of CTMP-SV40T, replacing the mCherry:*pac* fusion with *pac* the coding sequence only. The resulting plasmid was named CTP-SV40T. We designed a second expression cassette, consisting of the upstream region of *C. limacisporum* actin gene (Clim_evm35s218) and the SV40 terminator, separated by a new multiple cloning site. We used primers 14 and 15 to amplify the actin promoter from CAMP plasmid, and primers 16 and 17 to amplify SV40T from pSBtet. Primer 16 includes the restriction sites for the new multiple cloning site. Primers 15 and 16 contain an overlap region of 20 bases which were used to fuse the two fragments together into CTP-SV40T, linearized at the *NaeI* site, in a single Gibson Assembly reaction (New England Biolabs). The resulting plasmid was named pClimDC. All the cloning steps were controlled by Sanger sequencing.

### Construction of subcellular markers

pClimDC-NMM:tdT was constructed by fusing an N-myristoylation motif (NMM) to tdTomato. NMM was predicted on the *C. limacisporum* Src orthologue (Clim_evm93s153) using ‘NMT – The MYR Predictor’ online software (http://mendel.imp.ac.at/myristate/SUPLpredictor.htm) (Maurer-Stroh et al., 2002). The sequence coding for the predicted NMM sequence (GGCCSTETQQRQQPVRV) and a 21 bases region, starting from the second codon of tdTomato were included in primer 18. Primers 18 and 19 were used to build an NMM:tdTomato gene fusion by PCR using the pSM678 vector (a generous gift from Omaya Dudin) (Martin and Berthelot-Grosjean, 2009; Dudin et al., 2015) as a template. The resulting PCR product was cloned into pCR-Blunt II TOPO vector (Invitrogen), and checked by sequencing. The gene fusion was then amplified using primers 20 and 21 and cloned into pClimDC linearized at the *SalI* restriction site using Gibson Assembly (New England Biolabs). To build pClimDC-Lifeact:mCh, primers 22 and 23 were used to amplify the Lifeact:mCherry gene fusion from pONSY-Lifeact:mCherry (Parra-Acero et al., 2018). The resulting DNA fragment was inserted into the *SalI* site of pClimDC using Gibson Assembly (New England Biolabs). pClimDC-H2B:mV was constructed by fusing *C. limacisporum* histone 2B gene (Clim_evm20s1) with the coding sequence of the venus fluorescent protein. Primers 24 and 25 were used to amplify H2B gene from gDNA of *C. limacisporum* and inserted into the temporal CTV plasmid into *NotI* and *EcoRV* site that was constructed by inserting mVenus into pCR2.1Δf1/Kn^R^ (primers 26 and 27) with tubulin promoter and 3’UTR region of the *A. whisleri* actin gene (Awhi_g50). This insertion resulted in H2B:mVenus fusion. The gene fusion was then amplified using primers 28 and 29 and cloned into the *SalI* and *NheI* sites of pClimDC.

### Transfection by nucleofection

Transfections were performed using Amaxa^®^ 4D-Nucleofector^®^ and Primary Cell 4D-Nucleofector^®^ X Kit. Two days before transfection, *C. limacisporum* cells were seeded in 25 cm^2^ flasks (Falcon^®^ VWR, #734-0044) with 5 ml of marine broth. The day of transfection cells were counted using a Neubauer chamber. A total of 1.5 x10^6^ cells were used in each transfection experiment. Cells were harvested by centrifugation at 2500 *xg* for 5 minutes and the supernatant was removed completely. The pelleted cells were washed once with 100 μl ice-cold 1XPBS buffer (Sigma, #41639). During the centrifugation time of the washing step, the DNA mixture was prepared as follows: 3.3 pmol of plasmid at a concentration highly enough to assure that the DNA volume do not exceed 15% of the final reaction volume, together with 12.35 pmol of carrier DNA (pUC19) per condition. DNA mixture was added up to 24 μl chilled P3 4D-Nucleofector^®^ X Solution with previously added Supplement 1. The DNA solution was mixed thoroughly with the cells and a total volume of approximately 25 μl was transferred to a Nucleocuvette^®^ Stripe well. Nucleocuvette^®^ Stripes were gently tapped to ensure the cell suspension getting down to the bottom. Stripes were inserted into the 4D-Nucleofector^®^ X Unit and the EN-138 program was applied. After applying the pulse, 80 μl of MB medium were immediately added to the well. Mixed by pipetting and transferred to a 12 - well culture plate (Thermo Scientific Nunclon Delta Surface #150628), previously filled with 1 mL of MB medium. The plate was incubated at 23 ºC. The transfected cells could be observed after 24 - 48h using a fluorescence microscope.

### Enriching of resistant cells & clonal isolation

Two days after transfection, the cell culture was supplemented with puromycin at a concentration of 300 μg/mL and incubated for another 72 h. Cells were spun down in order to remove 800 μl of medium. Concentrated cells were spread on the surface of a marine agar Petri dish supplemented with 300 μg/ml puromycin, and after approximately seven to ten days single colonies were picked and amplified in liquid medium, always maintaining the drug pressure.

### Cryopreservation

Confluent culture of *C. limacisporum* - 500 μl, were frozen down in 500 μl of MB medium containing 10% DMSO (Sigma, #41639). Cryovials were transferred to Mr. Frosty™ Freezing Container and placed into a freezer to reach −80 ºC. Afterwards they were kept at −80 ºC for a long - term storage. In order to defrost and amplify the culture, 1 mL of frozen cells were transferred into a 25 cm^2^ culture flask filled with 9 ml of MB medium and reseeded the following day to reduce any putative adverse effect of DMSO on cells.

### Flow cytometry

The proportion of positively transfected cells and their distribution based on their fluorescent signal intensity were determined by flow cytometry at the Flow Cytometry Unit from Universitat Pompeu Fabra - Centre for Genomic Regulation, using a BD LSR Fortessa analyser (Becton Dickinson).

Transfection efficiency was analyzed 48 h post transfection (Fig. S2B, S2C) and strength of promoters (Fig. S2D) five days post transfection. In both cases samples were prepared as follow. Cells were scraped and harvested by centrifugation at 2000 *xg* for 5 minutes. After removing the supernatant, the pelleted cells were fixed resuspended in 500 μl of 4 % formaldehyde (Sigma-Aldrich, #F8775) in 1XPBS for 15 minutes. Afterwards cells were pelleted again by centrifugation and washed with 300 μl 1XPBS. All the steps were performed at the room temperature. Samples were stored at 4 °C until being processed.

BD FACSDiva™ software was used for data analysis and FlowJo™(v.10.7) for the data representation. SSC-A and FSC-A parameters were used to determine the cell population - P1. FSC-H and FSC-A were used to discriminate single cells from doublets - P2. Approximately 10.000 events were recorded from P2 population for determining mCherry positive population (P+). P+ were detected using a 561 nm laser with a 610/20 nm bandpass filter (red fluorescence) and differentiated from autofluorescence with a 780/60 nm bandpass filter. To set up gating and discriminate autofluorescence from the fluorescent signal of the positive population, cells were transfected with carrier DNA (pUC19), without the expression construct.

### Real-time quantitative PCR analyses

The number of plasmid copies integrated into the genome of different *C. limacisporum* transfected lines were estimated using real-time qPCR (Fig. S2D). *C. limacisporum* cultures were transfected with either CAMP, CTMP or carrier DNA (pUC19) only. After five days of growth, in the presence of 300 μg of puromycin, cells were harvested and genomic DNA (gDNA) was isolated. gDNA samples were diluted to a final concentration of 1 ng/μl.

The number of integrated plasmid copies in the different lines were quantified using iTaq Universal SYBR Green Supermix (Bio-rad #172-5121) in an iQ cycler and iQ5 Multi-color detection system (Bio-Rad). Primers that were used are indicated in the supplementary Table 2 (primers for actin promoter: 30, 31 and primers for tubulin promoter 9, 10). The total reaction volume was 20 μl. All reactions were run in triplicate. The program used for amplification was: (i) 95 ºC for 3 min; (ii) 95 ºC for 10 s; (iii) 60 ºC for 30 s; and (iv) repeat steps (ii) and (iii) for 40 cycles. Real-time data was acquired through the iQ5 optical system software v.2.1 (Bio-Rad). Plasmid copy numbers were estimated using the 2^−ΔΔCT^ method (Livak and Schmittgen, 2001) and are expressed as relative to the number of copies of genomic loci chosen at random (primers 32 and 32), whose proportion is expected to be constant between wild type and transfected lines.

### Stability of the transfected *C. limacisporum* lines

In order to evaluate the potential of these tools for downstream applications, we studied the stability of the transfection. We established five clonal lines (CAMP-transfected) starting from single colony isolates, – three from the *India* strain (I-1, I-2, I-3) and two from the *Hawaii* strain (H-1, H-2) and further analysed them. We first inquired about their stability over time. To test this, we started parallel liquid cultures from each clonal line, with and without selective pressure. The mean level of fluorescence was measured over three weeks, which corresponds to approximately 80 generations. When maintained in selective medium (+ puromycin), the level of mean fluorescence was constant for all the lines. This indicates transgenerational stability (Fig. S1B). However, when the same lines were cultured in non-selective medium (- puromycin), two out of the five lines showed a gradual decrease of fluorescent signal reaching the level of the negative control (untransfected cells) in less than two weeks (Fig. S1B). Negative cells were sorted and cultivated again in selective medium, and, interestingly, after seven days, the fluorescence signal was recovered (data not shown). The recovered fluorescence is in accordance with the presence of integrated plasmid copies, which a preliminary analysis of *nanopore* sequencing seems to verify (*Ocaña et al. in preparation*). Therefore, we would like to stress the need for testing the obtained transformants through several generations to select only the lines that maintain the expression of the fluorescent marker.

We then asked whether plasmid integration impacts the growth of *C. limacisporum*. For this, we determined the growth curves for two transformed lines and compare them with their corresponding wild-type strain. Results showed that exponential growth phase lasted 12 - 70h for wild-type *India* and *Hawaii* as well as for the I-3 transgenic line after which they reached the stationary phase. The H-1 transgenic line exhibited a slower growth rate but eventually reached the same number of cells at 160h (Fig. S1A). For this reason, we recommend to perform a growth test before selecting a line. Nevertheless, even though there was a decrease in growth in one of the lines as compared to the wild type, experiments with all lines were successfully carried out in parallel.

### Live cell imaging

For live cell imaging 3 x10^5^ cells were transferred to μ-Slide 4-well glass-bottom dish (Ibidi# 80426) containing 700 μl of marine broth and incubated for 2-4 hours before observation. Low melting agarose at a final concentration of 0.2% was added to reduce cell motility. Confocal microscopy of transfected stable lines with plasma membrane and / or nuclear marker in Fig. 3C and Fig. 4C was performed using Leica TCS SP8 STED 3X super-resolution microscope with an objective 100x/1.40 OIL STED WHITE and Leica TCS SP5 AOBS, with an HC PL APO 63x/1.40 Oil CS2 oil objective, respectively. Images in Figure 4A (pictures of cells with 1 and 4 nuclei), Figure 5A and Videos 1 - 3 were obtained using Andor iQ3 Spinning Disc microscope with an objective Nikon 100x/1.45 OIL TIRF. All remaining images and Movie 4 were performed with wide - field microscopy using a Zeiss Axio Observer Z.Q epifluorescence inverted microscope equipped with LED illumination and a Axiocam mono camera.

### Pulse-chase experiments

An aliquot of 1 mL from an exponentially growing culture of *C. limacisporum* was incubated for 30 min with the addition of 10 μl of a 1 mg/ml stock solution of FM4-64 (pulse). After incubation, the cell suspension was collected by centrifugation at 3500 x*g* - 1’, washed with 1 ml of marine broth (MB), centrifuged again and finally resuspended in 1 ml MB. The cells were incubated at 23 ºC for 15 min (chase) before mounting for microscopic observation.

### Cell fixation and phalloidin staining

*C. limacisporum* grown cell culture (1 mL) was harvested and spun down at 2500 *xg* and washed with 1XPBS. The cells were resuspended in a fixative - 4% formaldehyde for 15’ at the room temperature (RT) and afterwards washed 3 times with 1XPBS. For actin staining in Fig. S4, 1 μl of Alexa Fluor™ 488 Phalloidin (Thermo Fisher Scientific #21838) was added to 10 μl of concentrated cell sample and incubated for 15’ at RT in the darkness. Pictures were taken with the wide-field microscopy (see above).

### Image Analysis

Image analysis was performed using ImageJ software (version 2.0)(Schneider et al., 2012). Cells diameter in Fig. S6 and Fig. 5 was calculated from perimeter measurements obtained manually with an oval selection tool. All Figures were assembled with Inkescape (version 1.0).

## SUPPLEMENTARY FIGURES

**Figure S1.**
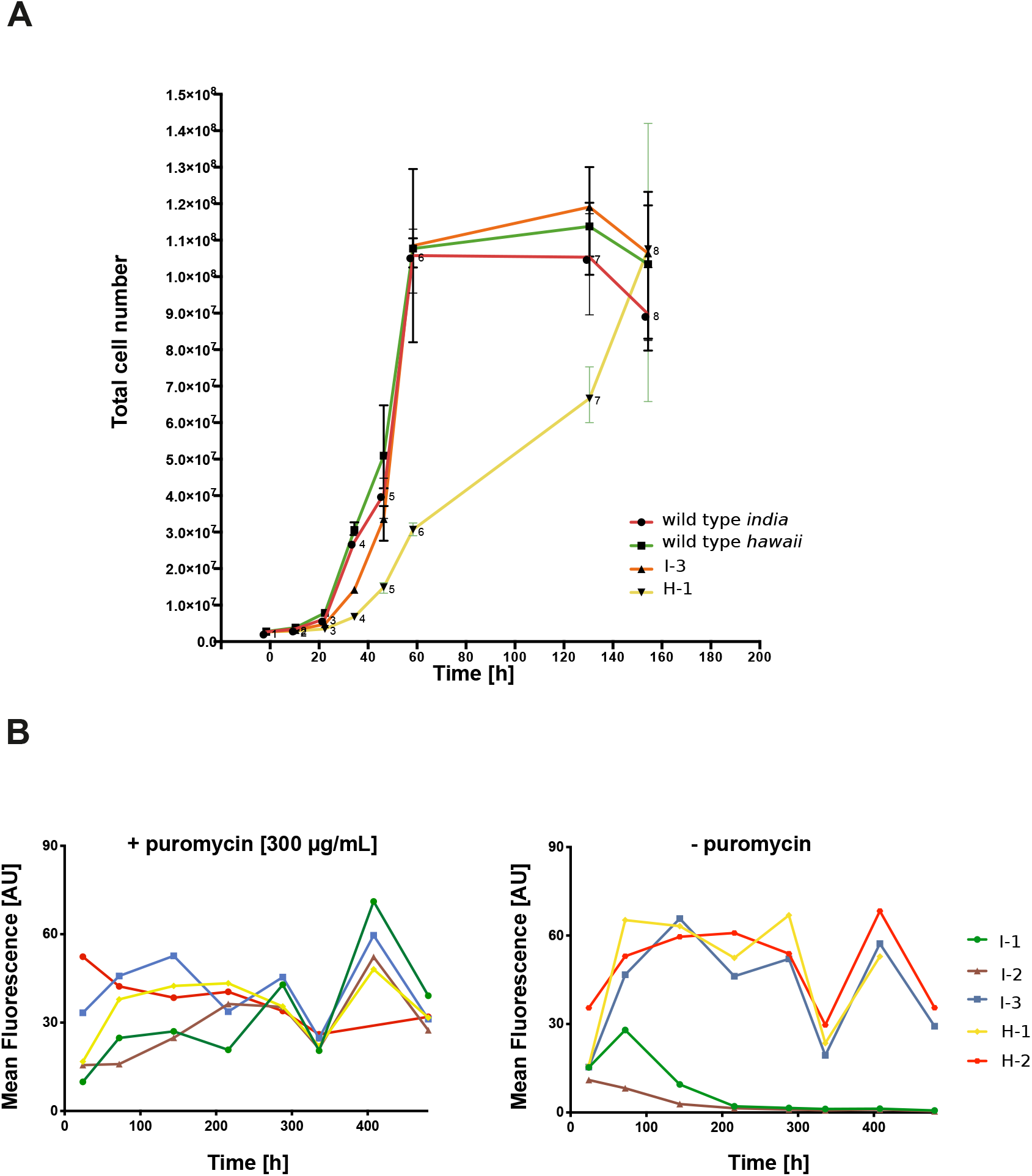
Characterization of transfection lines. **A.** Growth curves of wild type *C. limacisporum* strains: *India* and *Hawaii* and corresponding transfected lines. Each time point is generated by the average of the triplicates from manual cell counting and plotted on a linear scale. **B.** Testing stability of transformants overtime in the presence or absence of selective pressure. Mean fluorescence of the mCherry protein in the CAMP - transfected *C. limacisporum* culture was measured over time. Two parallel liquid cultures coming from a corresponding colony isolate, were maintained with and without selective pressure for 16 days. In the selective medium all five lines expressed the constant level of fluorescence whereas in non-selective medium two out of five lines (I-1 and I-2), showed a gradual decrease of fluorescent signal eventually reaching the equivalence of the negative control.

**Figure S2.**
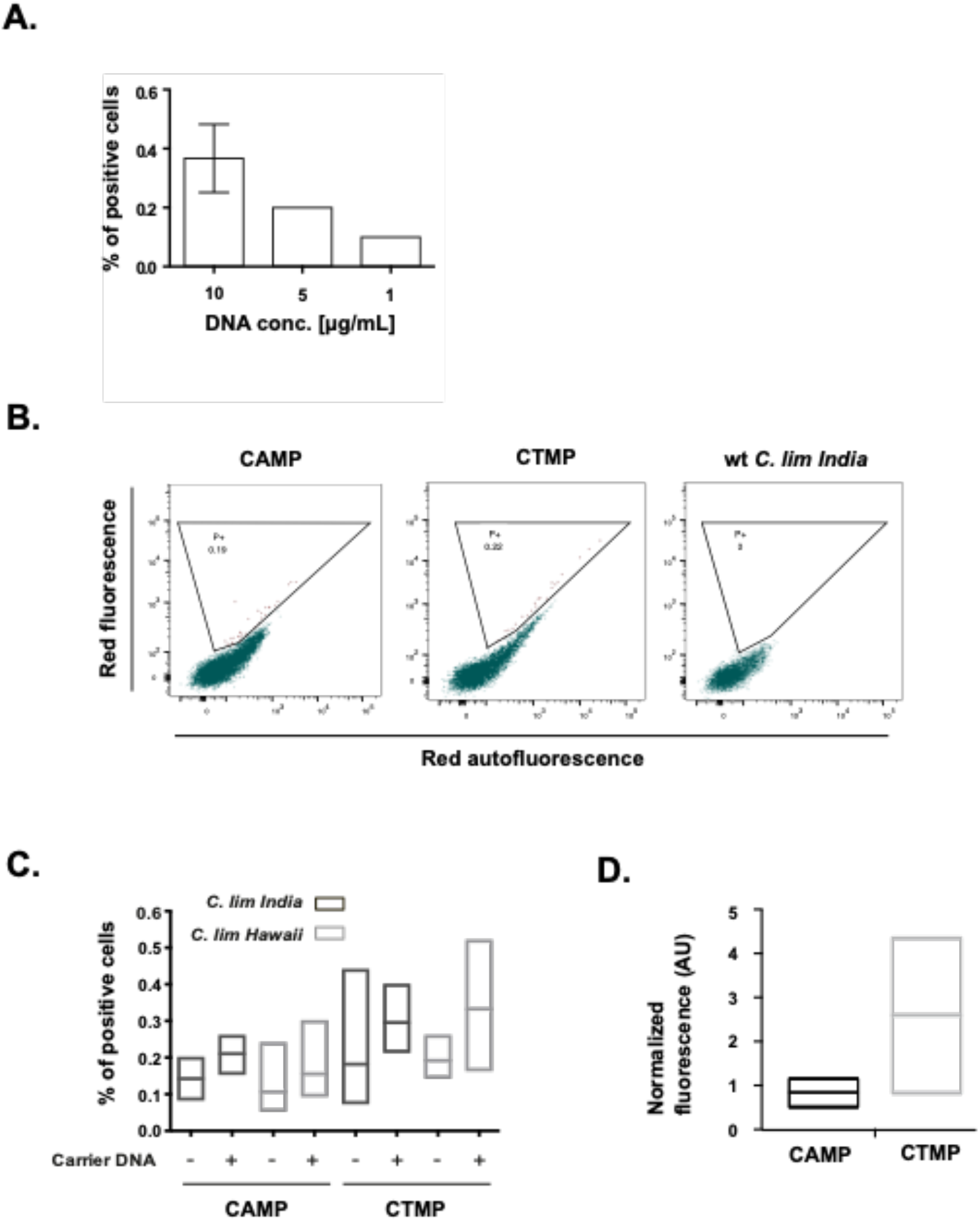
Characterization of transfection tools. **A.** Number of positively transfected cells (average number from a triplicate sample) in relation to different plasmid DNA concentrations (transfection construct: CTMP). **B**. Representative flow cytometry distribution of *C. limacisporum India* cells transfected with CAMP and CTMP plasmids, and untransfected cells. Area selected (P+) represents the mCherry positive population, which was defined in relation to untransfected population. **C.** Percentage of positive cells obtained by transfecting both promoter variants (actin promoter - CAMP and tubulin promoter - CTMP) in the presence or absence of the carrier DNA (pUC19), for both *C. limacisporum* strains, *India* and *Hawaii*. **D.** Calculation of the strength of each promoter variant (CAMP vs CTMP). The mean fluorescence intensity of mCherry fluorescent protein was measured by flow cytometry and normalized with plasmid copy number quantified with RT-qPCR. Transfected strain: *C. limacisporum India*.

**Figure S3.**
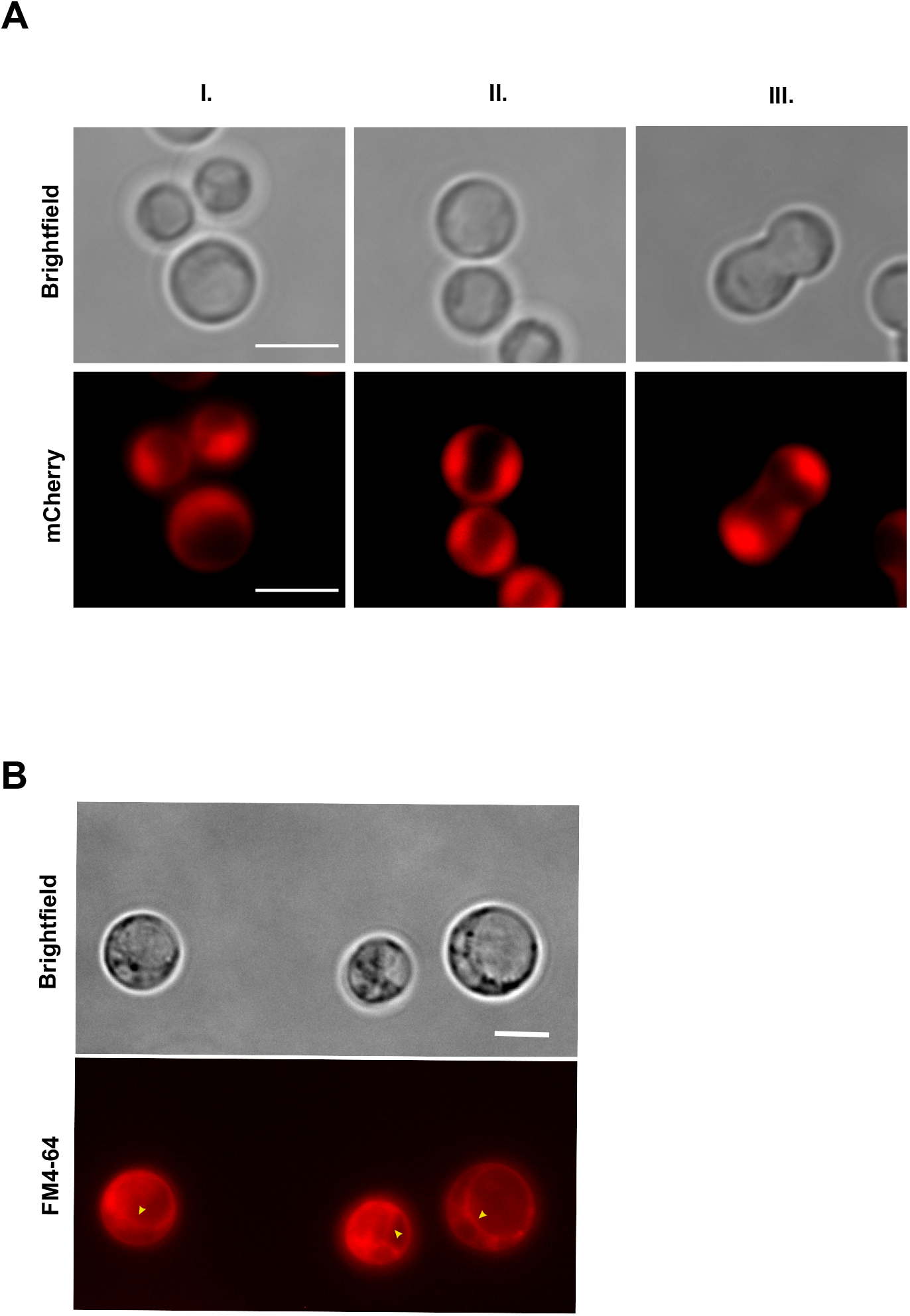
**A. Cytoplasm visualization of *C. limacisporum*.** Cells were transfected with the CAMP construct revealing a ‘crescent-like’ shape cytoplasm that is present on one side (I.), or on both sides (II.) of the vacuole and, during cell division (III.). Cells were imaged using widefield fluorescent microscopy. **B. Testing for the presence of a vacuole in *C. limacisporum*.** Vital staining with the lipophilic membrane dye FM4-64 evidenced *C. limacisporum* vacuoles (arrowheads).

**Figure S4.**
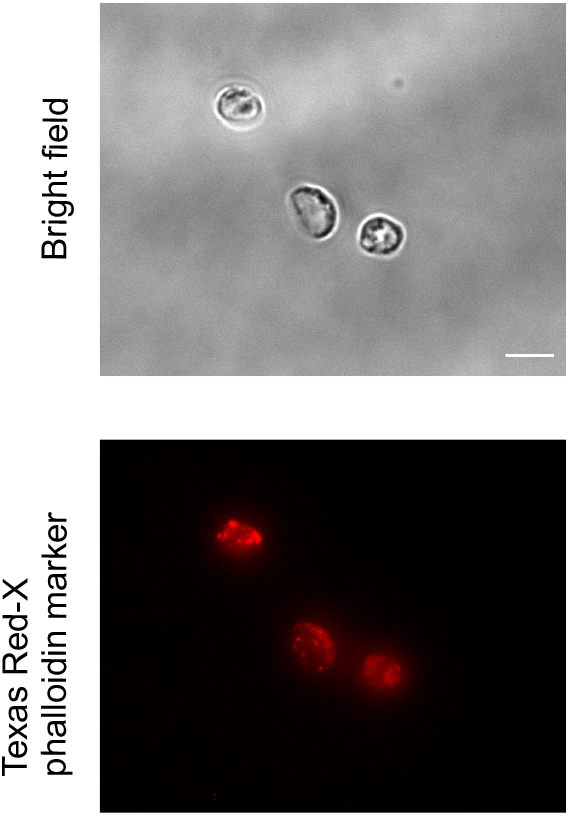
Actin filaments staining in *C. limacisporum* on fixed cells. Fixed cells were stained with Texas Red-X phalloidin marker (10 μm) to validate results obtained from cell transfection with F-actin marker, pClimDC-Lifeact:mCh (Fig. 3C). Cells were imaged using widefield fluorescent microscopy.

**Figure S5.**
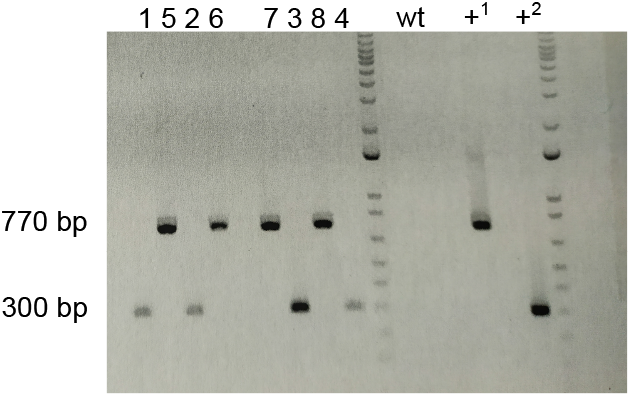
Amplification of transfection constructs (pClimDC-H2B:mV and pClimDC-Lifeact:mCh) from gDNA of stably transfected lines. Positive amplification of the corresponding transfected plasmid in transformed lines, confirm plasmid integration into the genome. Four independent cultures of each transformed lines were initiated from a single colony, and used for gDNA isolation. pClimDC-H2B:mV: transformed line 1 – 4, pClimDC-Lifeact:mCh transformed line: 5 – 8. wt = wild type (untransfected line), +^1^ = pClimDC-H2B:mV plasmid, +^2^ = pClimDC-Lifeact:mCh plasmid.

**Figure S6.**
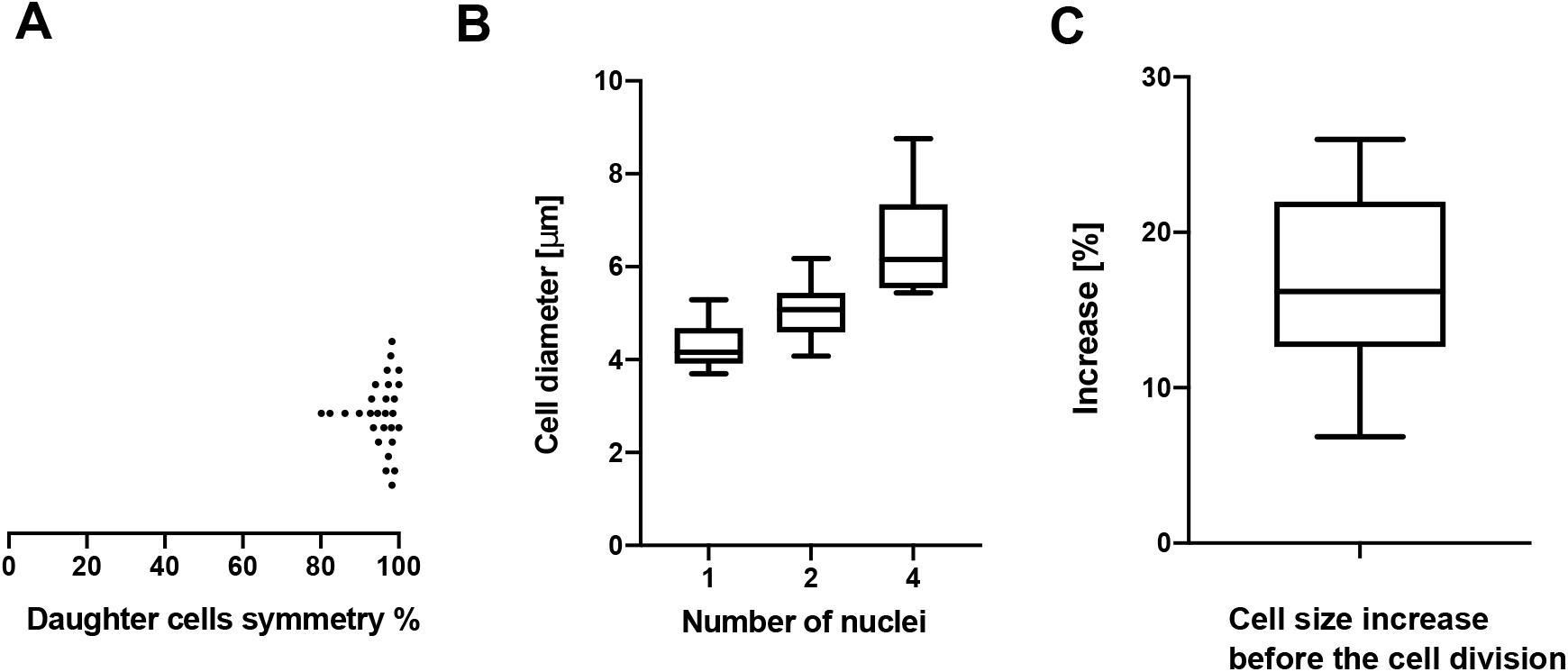
**A.** Symmetry of daughter cells after cell division. Each point represents % of similarity between the diameters of two daughter cells. In total 58 measurements were taken (28 pairs). **B.** Cell size in relation to number of nuclei. Number of nuclei was visualized with the nuclear marker (pClimDC-H2B:mV). N° of measurements: mononucleate cells: 20, binucleate cells: 30, > 2 nuclei cells: 10. **C**. Cell size increase before a cell division. Diameter of each cell was measured just after a cell was born from a mother cell and before the next cell division. Number of measurements: 25. Cell diameters were calculated with ImageJ software.

**Figure S7.**
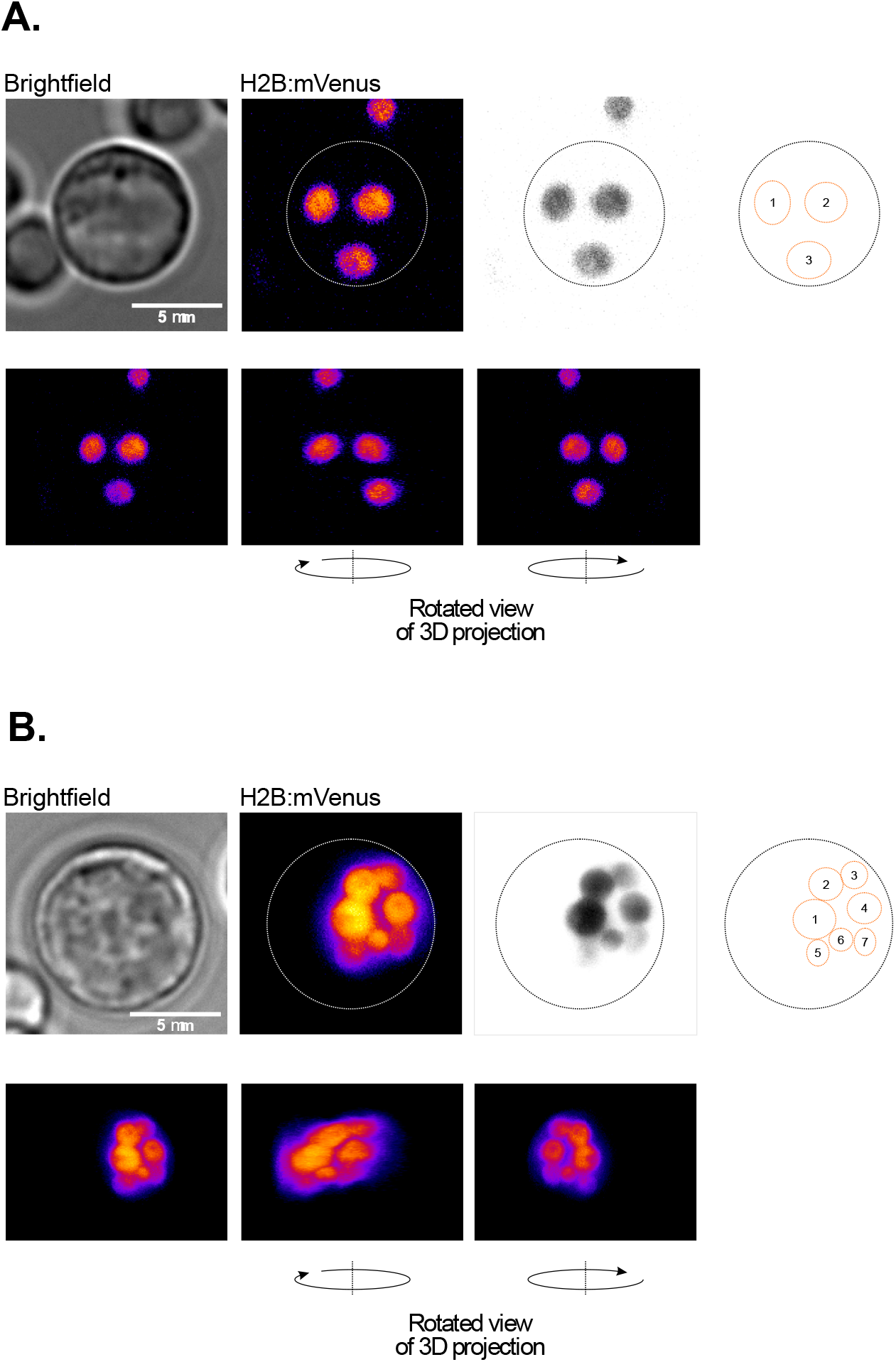
*C. limacisporum* coenocytic cells with odd number of nuclei. Visualization of nuclei in *C. limacisporum Hawaii* coenocytic cells. Images show cells with **A.** three and **B.** seven nuclei with their respective sketches facilitating nuclei counting. Cells were imaged with wide-field fluorescent microscopy. 3D projection was performed with ImageJ. Scale bar 5μm.

**Suppl. Table 1.** List of antibiotics and herbicides tested on *C. limacisporum*. Puromicyn was chosen as the most optimal drug taken into account the time needed to promote cell death in a given population. The most efficient concentration was 300 μg/ml. Drug concentrations tested: 50 and 100 μg/ml in the indicated solvent by manufacturer unless indicated otherwise. Different drug concentrations tested: Puromycin: 100 - 500 μg/ml, Benomyl: 20 - 500 μg/ml, Carboxine: 20 - 300 μg/ml, Nourseothricin: 10 - 50 μg/ml, Fluoretic Acid: 250 - 30 μg/ml.

| Name of a drug | Effect: Strong (++), mild (+), none (-) |
| --- | --- |
| Puromycin | ++ |
| Carboxine | + |
| Nourseothricin | + |
| Phleomycin | + |
| Benomyl | + |
| Glusifonato | + |
| Genetycin (G418) | + |
| Fluoretic Acid | - |
| Hygromycin | - |
| Fungizone | - |
| Spectomycin | - |
| Paclitaxel | - |
| Chloramphenicol | - |
| Gentamicin | - |
| Kanamycin | - |
| Methotexate | - |
| Penicilin+Streptomycin | - |
| Glyphosate | - |

**Suppl. Table 2.**
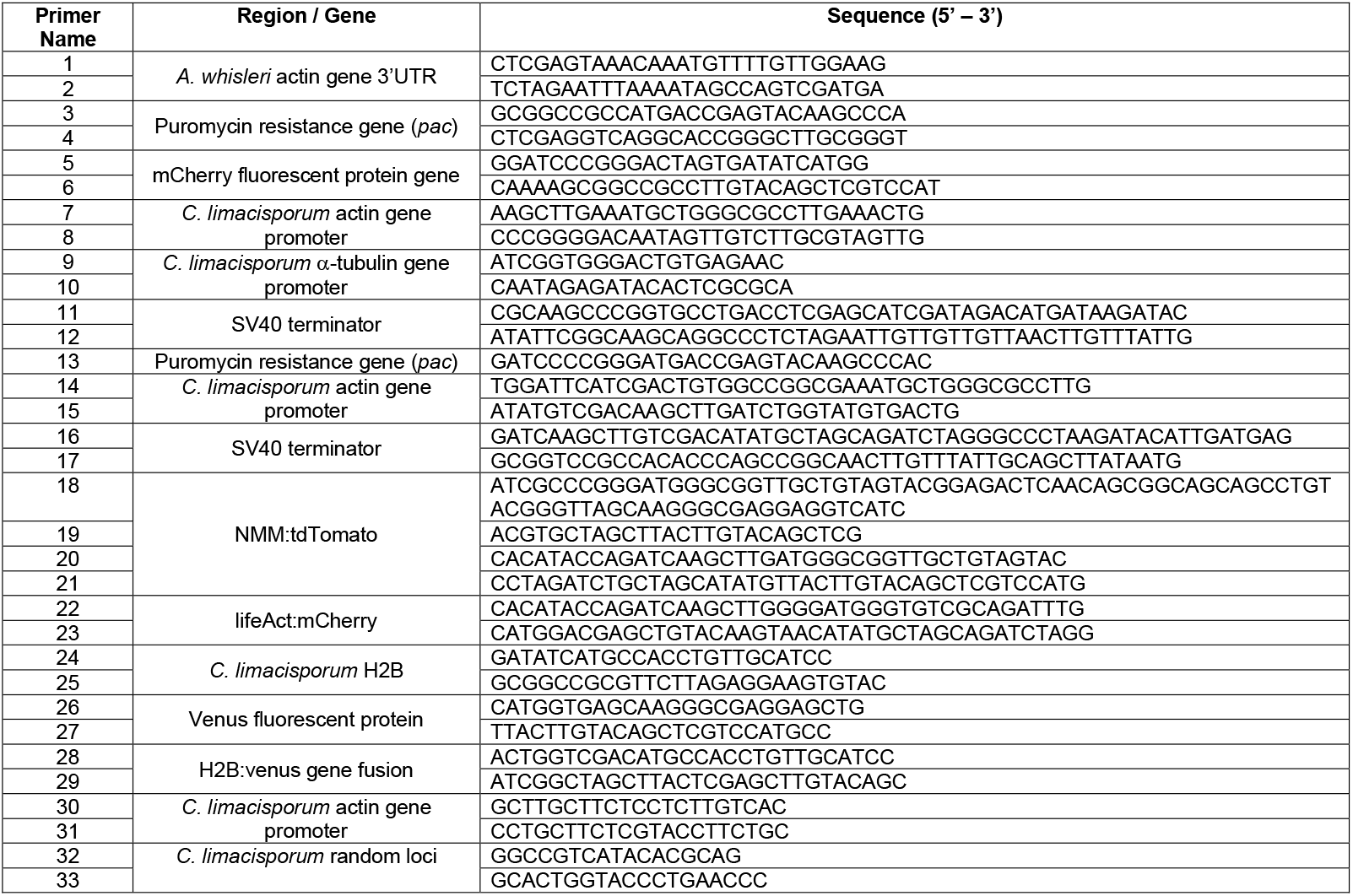
List of primers used to build *C. limacisporum* expression vectors.

## SUPPLEMENTARY MOVIES

**Movie 1.**

Time-lapse of *C. limacisporum Hawaii* cells transfected with the plasma membrane marker (pClimDC-NMM:tdT) and the nuclear marker (pClimDC-H2B:mV). Images were taken every 5 minutes during 23h with a spinning disc microscope. The movie shows binary fission of *C. limacisporum* cells as well visualization of nuclei division. The images from the movie were used to represent the decoupling of cell division from nuclear division in the Figure 5A. Scale bar represents 5 μm.

**Movie 2.**

Time-lapse of *C. limacisporum Hawaii* cells transfected with the plasma membrane marker (pClimDC-NMM:tdT) and the nuclear marker (pClimDC-H2B:mV). Images were taken, every 5 minutes during 23h with a spinning disc microscope. The movie presents: quadrinucleate coenocyte that creates a quatrefoil-like shape (*arrow 1*); a bi-nucleate cell that forms two lobes, after which returns to a spherical shape and continues to double its nuclei, after which forms a quatrefoil-like shape and divides leading to the rise of four mononucleate cells (*arrow 2*); a quatrefoil-like shape cell (*arrow 3*). Scale bar represents 5 μm.

**Movie 3.**

Time - lapse of *C. limacisporum Hawaii* cells transfected with the plasma membrane marker (pClimDC-NMM:tdT) and the nuclear marker (pClimDC-H2B:mV). Images were taken every 5 minutes during 23h with a spinning disc microscope. The movie presents binary fission of cells (*arrow 4*), as well as a bi-nucleate cell becoming a coenocyte (*arrow 5*) followed by the release of new cells in a budding - like manner. Scale bar represents 5 μm.

**Movie 4.**

Time-lapse of a wild type *C. limacisporum* coenocyte cell and the release of amoebas. Images were taken with a wide-field optical microscopy. Scale bar represents 5 μm.

